# Diet-Driven Gut Microbiome Differences Between Cohabiting Han and Yi Ethnic Groups in Liangshan, China

**DOI:** 10.64898/2026.01.06.698051

**Authors:** Chenglu Jiang, Bin Hu, Cheng Yang, Xiaoli Yuan, Maoping He, Xingyi Yan, Yuzhu Li, Guo Li, Donglin Tan

## Abstract

**Background:** While ethnic variations in gut microbiota are well-documented, their persistence in genetically distinct populations sharing the same geographic environment remains poorly understood. This study investigates the Han and Yi ethnic groups cohabiting rural Southwest China, where contrasting dietary traditions persist under identical environmental conditions.

**Methods:** We analyzed fecal samples from 100 healthy adults (50 Yi/50 Han) using 16S rRNA gene sequencing (V3-V4 region). Dietary intake was quantified via three 24-hour recalls validated with plasma biomarkers. Multivariate analyses (DESeq2, PERMANOVA) controlled for age, sex, and BMI covariates.

**Results:** Yi individuals exhibited significantly higher Prevotellaceae (log2FC=1.82, padj=0.028) and Succinivibrionaceae (log2FC=2.15, padj=0.013) abundances, corresponding to their buckwheat-rich diet (58.5 ± 12.3% vs Han 3.3±1.8%, p<0.001)

Han microbiota showed enriched Bacteroidaceae (log2FC=1.24, padj=0.041), associated with higher animal fat intake (23.6% vs Yi 1.8%, p<0.001)

Dietary factors (buckwheat/animal fat intake) explained greater microbiota variation than ethnic genetic ancestry (PERMANOVA R²=0.10 vs 0.07, p < 0.05)

**Conclusion:** Cohabiting Han and Yi populations maintain distinct gut microbiota compositions primarily driven by dietary differences. These findings highlight diet as a stronger determinant than genetic ancestry in shaping microbial communities under shared environments. Future studies should validate these observations with metagenomic sequencing.

## Introduction

The gut microbiome constitutes a complex ecosystem that plays fundamental roles in host metabolism and health maintenance[1; 2]. Accumulating evidence identifies dietary patterns as the primary non-genetic determinant of microbial community structure [3; 4], with long-term dietary habits exerting profound effects on microbial composition and metabolic output[5]. However, the impact of ethnic-specific dietary traditions—particularly those adapted to unique environmental pressures—on gut microbiota configuration and host health remains underexplored[6]. Current understanding derives primarily from studies comparing geographically isolated populations[7; 8], which inherently conflate genetic and environmental influences. While large-scale surveys demonstrate joint contributions of ethnicity and geography to microbiome variation[9; 10] , few studies have examined ethnically distinct groups coexisting within the same microenvironment. This limitation obscures our understanding of how dietary practices per se drive microbial divergence when environmental confounders are minimized.

Notably, indigenous populations maintaining traditional diets exhibit microbial signatures distinct from urbanized groups[11; 12]. For instance, high-fiber diets characteristic of hunter-gatherer populations select for taxa specialized in complex carbohydrate degradation[8] . Functional redundancy—the coexistence of phylogenetically distinct taxa performing similar metabolic roles—has been proposed as a key ecological mechanism conferring microbiome stability[13; 14]. However, the role of such adaptations in sustaining ethnic-specific microbial configurations remains poorly understood[15].

- The Yi and Han populations of China’s Liangshan Prefecture present a unique natural experiment. Cohabiting the same rural communities at 1,500-2,500m altitude, these groups maintain starkly contrasting dietary traditions: Yi: Buckwheat-dominated (46.8% vs Han 2.2%), high-resistant starch diet
- Han: Rice/animal-fat based (45.3% vs Yi 2.5%), Western-influenced diet while sharing identical environmental exposures (climate, water sources, etc.).

This study leverages 16S rRNA gene sequencing and covariate-adjusted analyses to test three specific hypotheses:

1. Ethnic dietary differences drive microbiome divergence exceeding genetic influences
2. Yi’s buckwheat-rich diet selects for microbial consortia with enhanced functional redundancy
3. These microbial differences associate with measurable variations in host metabolic parameters

## Results

### Cohort Characterization and Dietary Profiling

Tis study recruited participants from Xichang, a city located in southwest of sichuan province, china. Fecal samples were obtained from 113 participants. After excluding participants with missing data, 100 participants were included for analyses (Supplementary Table1). Age, sex, BMI, ethnicity, and food type were selected as covariates according to previous research[16; 17]. The conceptual framework (Fig.1A) illustrates the analytical pathway for identifying ethnic-specific characteristics, in parallel to fecal sampling, we collected subject metadata including ethnicity, BMI, age, sex, blood biochemical measurements. Comparative analysis of demographic parameters (Fig.1B)revealed distinct ethnic profiles. Yi participants demonstrated significantly higher age values (p = 0.039) and body mass indices (p = 0.047) compared to Han counterparts. Gender distribution showed no statistical variation between groups, confirming effective matching for this potential confounder; these baseline differences were accounted for in subsequent microbiota analyses through covariate adjustment. Dietary patterns exhibited marked ethnic stratification(Fig.1C). Yi participants demonstrated substantially higher consumption of buckwheat (58.5% vs 3.3%) and Bacon (23.2% vs 0.4%) compared to Han counterparts. Conversely, Han individuals showed a greater intake of rice (45.3% vs 2.5%), vegetables (17.7% vs 10.9%), fresh meat (23.6% vs 1.8%), and fruits (9.7% vs 3.0%). Bacon and rice consumption patterns were comparable between groups.

**Figure 1.**
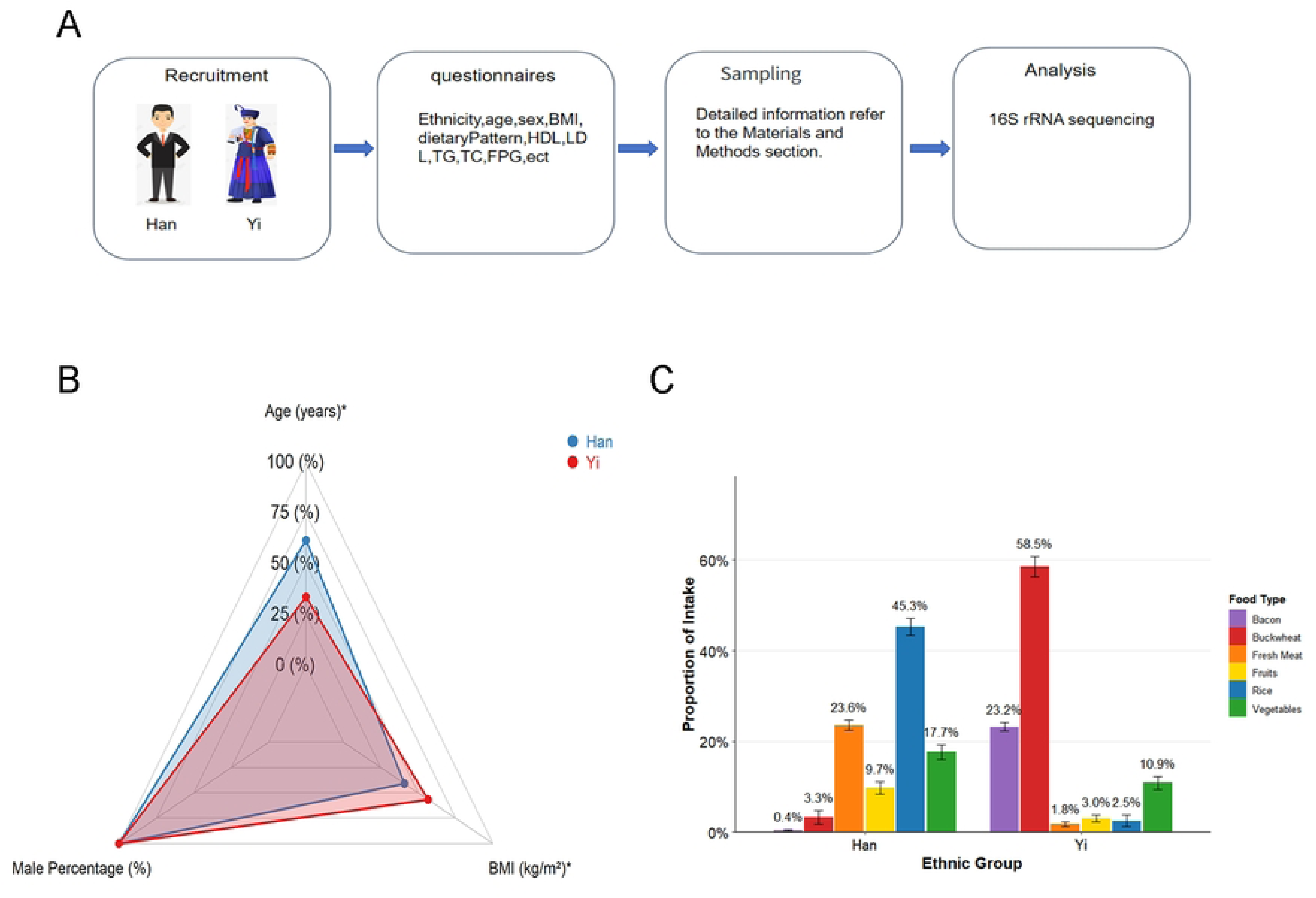
Cohort Design. **A.** An overview of study design and data collection regime, including volunteer recruitment, metadata questionnaire investigation, and dietary pattern investigation. **B.** The radar chart displays the mean values of age, male percentage, and body mass index (BMI) for the Yi (red) and Han (blue) groups. **C.** Proportional consumption of six major food categories is presented by ethnic group. Food types include bacon, buckwheat, fresh meat, fruits, rice, and vegetables. Analysis adjusted for age, sex, and BMI where applicable.

Dietary Factors Outweigh Ethnicity in Shaping Gut Microbiota Diversity, Community Structure, and Taxonomic Composition in Han-Yi Populations Shannon’s α Diversity Index[18], which weights both microbial community richness (observed operational taxonomic units [OTUs]) and evenness (Equitability), showed significant variation across ethnicities. In our study, microbial Shannon’s α Diversity (Fig.2A)demonstrated significant ethnic stratification (ANCOVA *p*=0.017, η²=0.06). Yi individuals exhibited consistently higher Shannon indices relative to Han counterparts (median: 4.2 vs 3.8; IQR: 3.9-4.5 vs 3.5-4.1). This pattern persisted after adjusting for age, BMI, and dietary covariates, suggesting intrinsic ethnic determinants of gut ecosystem complexity. Ethnicity explained 7% of microbiota variation (R² = 0.07, p = 0.046), while dietary factors collectively accounted for 10% of variation (R² = 0.10, p = 0.030). The primary separation axis (PCoA1, 38.9% variation) showed strong associations with specific dietary components, including rice, buckwheat, bacon, and fresh meat intake. This ethnic segregation pattern (PERMANOVA p < 0.05) highlights the combined influence of genetic and dietary factors on microbial community structure. Striking differences emerged at the family taxonomic level (Fig. 2C). The Yi microbiota exhibited elevated Prevotellaceae (19.3% vs. 8.7%, FDR = 0.001), representing their most abundant family and consistent with high-resistant starch intake. Concurrent Succinivibrionaceae enrichment was observed in Yi (4.2% vs 1.0%, FDR = 0.011), a taxon associated with complex carbohydrate metabolism. While significant enrichment of Bacteroidaceae in Han individuals (15.2% vs 7.0%, FDR=0.011) suggests enhanced bile acid metabolism, potentially reflecting habitual consumption of animal-derived foods. This enterotype feature aligns with Western-style diets where Bacteroidaceae serves as a primary converter of primary to secondary bile acids, with dual implications for lipid absorption and colorectal cancer risk[4; 6]. Quantitative comparison confirmed significant compositional differences (Fig. 2D). Prevotellaceae abundance was 2.2-fold higher in Yi versus Han, while Succinivibrionaceae showed 4.3-fold enrichment in Yi. Bacteroidaceae demonstrated reciprocal enrichment in Han. These differentially abundant families suggest distinct functional adaptations: Yi microbiota specialized in starch/carbohydrate processing (Prevotellaceae-dominant), while Han microbiota showed enhanced bile acid metabolism capacity (Bacteroidaceae-enriched).

**Figure 2.**
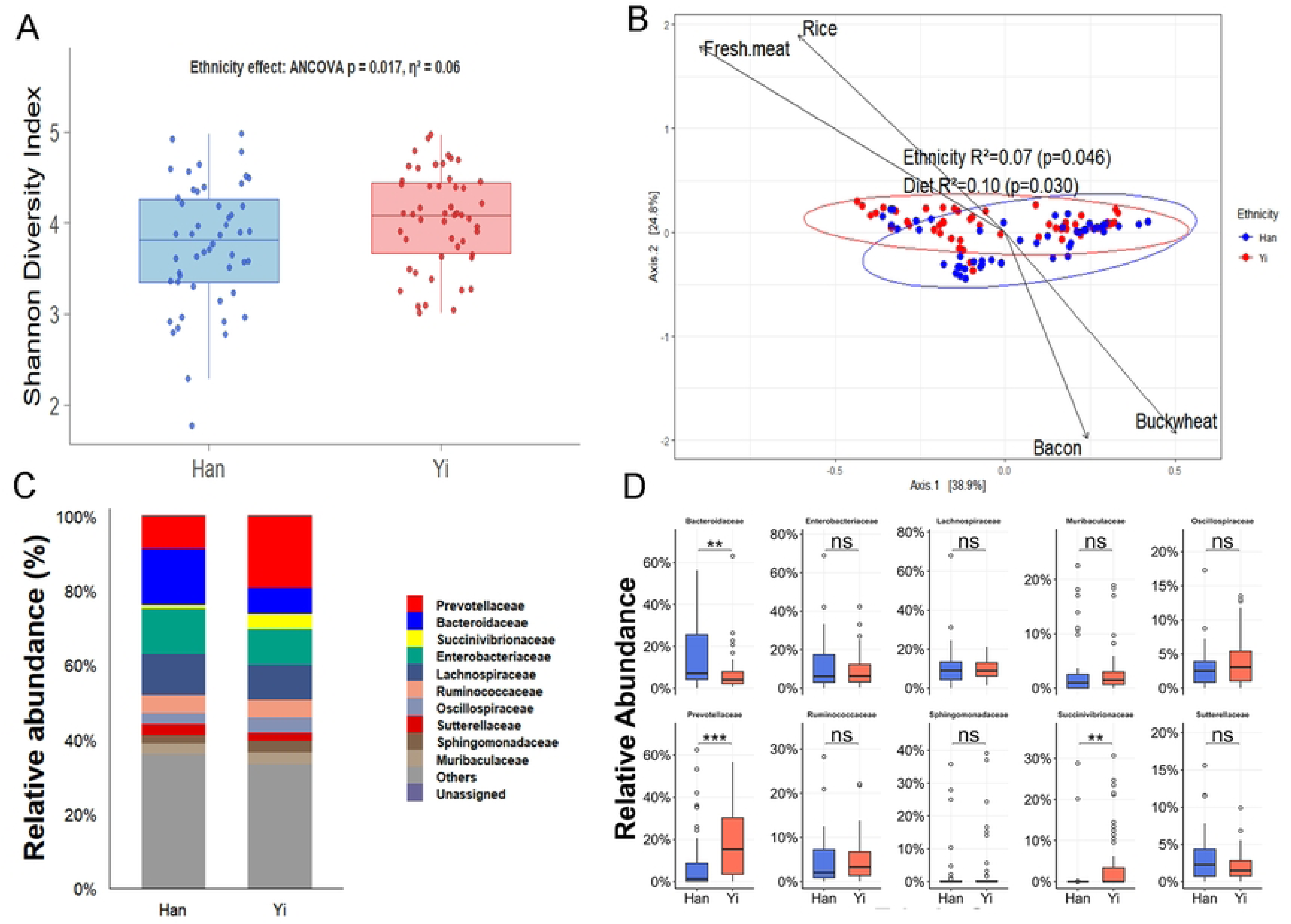
thnic and Dietary Drivers of Gut Microbiota Variation **A.**Comparative analysis of Shannon diversity indices between Han and Yi populations demonstrates significant ethnic effects (ANCOVA: p = 0.017, η² = 0.06). **B.** Ordination diagram demonstrating the relative contributions of ethnicity and dietary patterns to gut microbial community variation, ethnicity explains significant variance (R² = 0.07, p = 0.046), dietary factors account for greater explanatory power (R² = 0.10, p = 0.030). **C.** Comparative profiling reveals differential enrichment of 10 core bacterial families in Han and Yi gut microbiomes at family level. **D.** Ethnic disparity in 10 core bacterial family abundances reveals differential enrichment patterns between Han and Yi gut microbiomes (FDR-adjusted p < 0.05). Analysis adjusted for age, sex, and BMI where applicable.

### Phylogenetic and Ecological Divergence of Gut Microbiota Reflects Dietary Adaptation in Yi and Han Ethnic Groups

Building upon the compositional disparities observed in family-level abundances (Fig. 2D), we investigated the evolutionary and ecological relationships among differentially abundant families. Phylogenetic analysis revealed that Yi-enriched Prevotellaceae and Succinivibrionaceae formed a highly supported monophyletic clade (bootstrap = 92%, Fig. 3A), suggesting shared evolutionary trajectories potentially driven by adaptation to carbohydrate-rich diets. Conversely, Han-enriched Bacteroidaceae occupied a distant phylogenetic position, consistent with its specialized role in bile acid metabolism. The co-occurrence network (Fig. 3B), constructed using stringent thresholds (|Spearman ρ| > 0.3 and FDR-adjusted p < 0.05), revealed 21 significant interactions among 10 bacterial families. Microbial co-occurrence networks further demonstrated functional coherence within ethnic groups: Prevotellaceae and Succinivibrionaceae exhibited strong positive interactions (Spearman r > 0.6, FDR < 0.01) in Yi individuals (Fig. 3B), indicating synergistic roles in starch degradation. In contrast, Bacteroidaceae served as a hub for bile acid-transforming consortia in Han individuals, co-occurring with secondary bile acid producers (e.g., Lachnospiraceae). Principal coordinates analysis confirmed significant structural divergence between ethnic groups (PERMANOVA R² = 0.051, p = 0.002; Fig. 3C), though the modest effect size implies concurrent influences from non-genetic factors. Notably, the first principal component (PC1, 31.8% variance) separated Yi (negative scores) and Han (positive scores) microbiomes, aligning with the Prevotellaceae-driven versus Bacteroidaceae-driven community structures."

**Figure 3.**
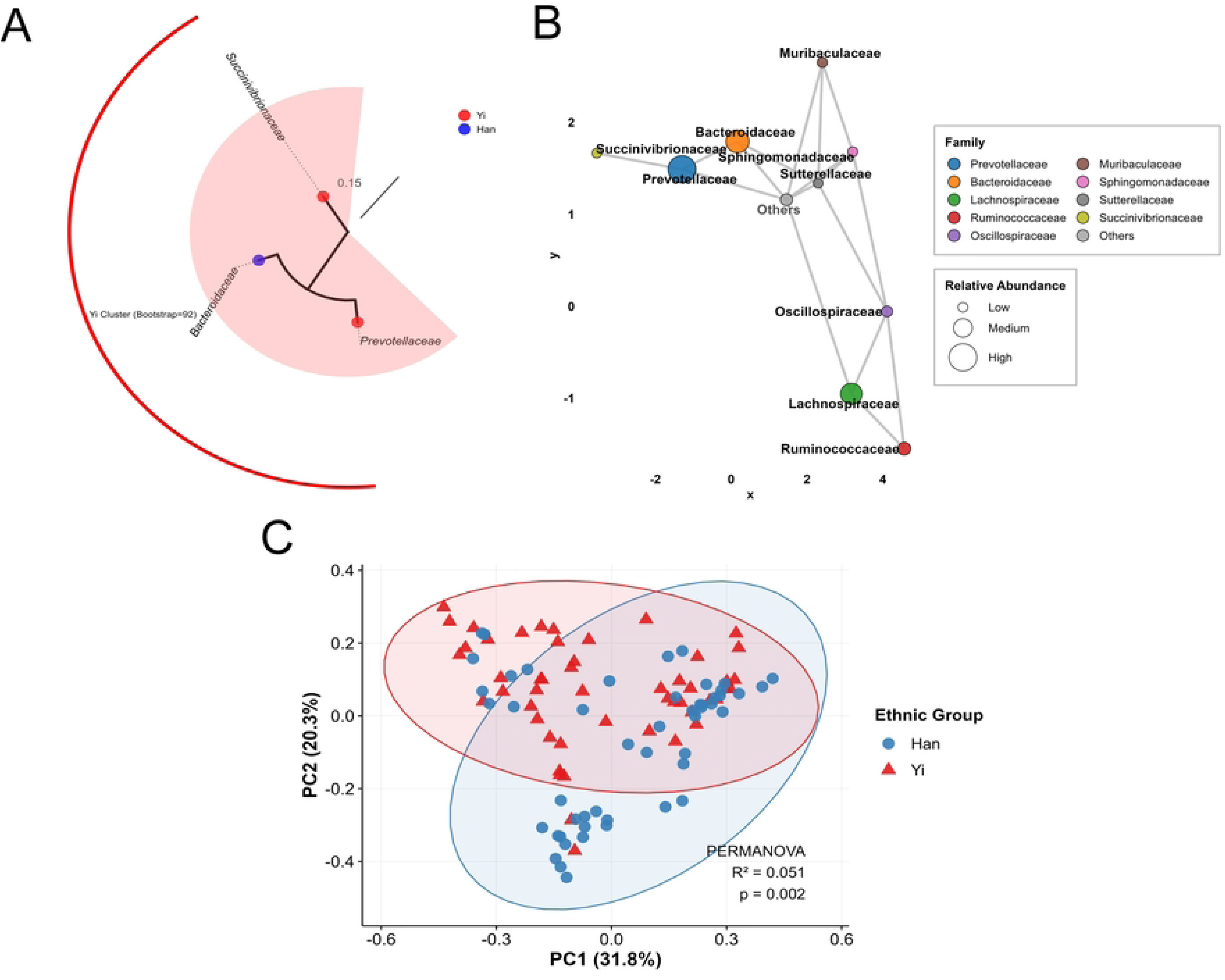
Evolutionary and ecological divergence of gut microbiomes between Yi and Han ethnic groups **A.** Evolutionary reconstruction of three dominant bacterial families reveals distinct clustering patterns associated with ethnic origin, Yi-derived strains form a strongly supported monophyletic clade (robustness validation > 90%), demonstrating substantial genetic divergence from Han counterparts. **B.** Ethnicity-driven restructuring of microbial interaction networks reveals fundamental topological reorganization of gut microbiota associations across ethnic cohorts. **C.** Beta diversity analysis demonstrates statistically robust segregation of Han and Yi gut microbiomes along primary ecological vectors (PERMANOVA p < 0.005). Analysis adjusted for age, sex, and BMI where applicable.

### Integrated Analysis of Diet-Driven Gut Microbiome Specialization and Ecological Adaptations

Our comprehensive analysis reveals three interconnected dimensions of gut microbiome adaptation to ethnic dietary patterns. First, functional specialization analysis (Fig. 4A) demonstrates distinct metabolic profiles between Yi and Han populations. Yi individuals, consuming high-buckwheat diets, exhibit significant enrichment in glycan metabolism pathways (1.17-fold, p=0.005) through their Prevotellaceae-dominated consortium, while Han microbiomes show elevated terpenoid metabolism (1.06-fold, p=0.006) via Bacteroidaceae-mediated bile acid transformation, reflecting adaptation to animal fat consumption.

**Figure 4.**
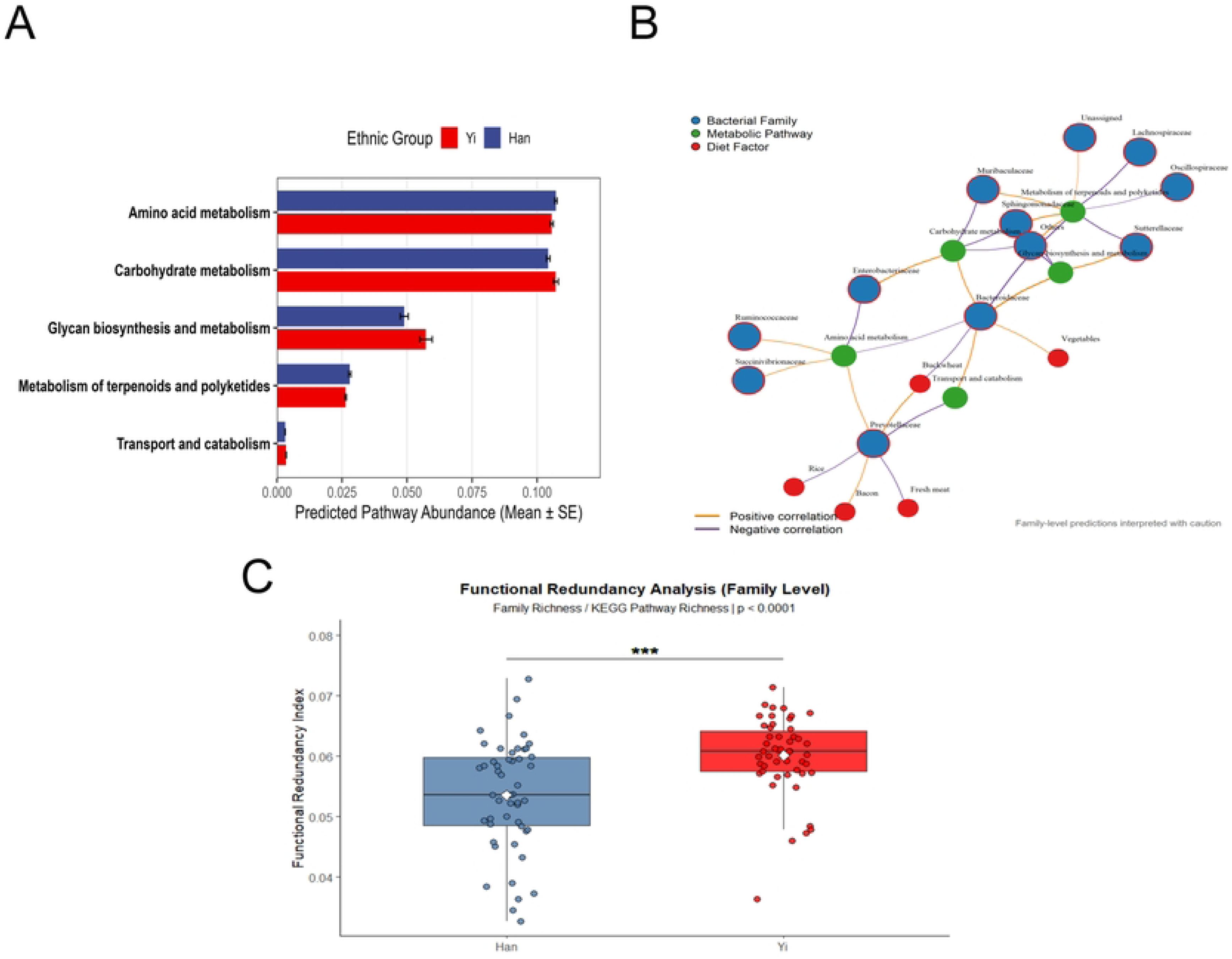
**A.** Functional profiling reveals divergent enrichment of core biochemical pathways between ethnic cohorts. **B.** Integrated network modeling reveals ethnic-constrained associations between dietary patterns, bacterial families, and metabolic pathways. **C.** Ethnic differences in functional redundancy index. Analysis adjusted for age, sex, and BMI where applicable.

The tripartite network analysis (Fig. 4B) elucidates key diet-microbe-function relationships, including: (1) the buckwheat-Prevotellaceae axis (r=0.42, p=1.1×10⁻⁵) with enhanced amino acid metabolism (r=0.41); (2) bacon consumption associations with specific bacterial families (e.g., Enterobacteriaceae); (3) Bacteroidaceae’s dual role in glycan biosynthesis (r=0.66) and terpenoid suppression (r=-0.58). These findings are complemented by functional redundancy assessment (Fig. 4C), showing Yi microbiomes possess superior resilience (12% higher redundancy, p<0.0001) through Prevotellaceae-Succinivibrionaceae networks (bootstrap=92%), albeit with adiposity risk (β=+41.8), while Han microbiomes demonstrate specialized bile acid metabolism with potential cardiovascular benefits.

This integrated view demonstrates how traditional diets architect functionally distinct gut ecosystems through: (1) pathway specialization (Fig. 4A), (2) network-level adaptations (Fig. 4B), and (3) ecological stability trade-offs (Fig. 4C), providing a framework for understanding diet-microbiome-health relationships across ethnic groups.

## Discussion

This comprehensive study elucidates how traditional dietary practices fundamentally reconfigure gut microbiome architecture and metabolic functionality in cohabiting Han and Yi populations in Liangshan, Southwest China, with profound implications for cardiometabolic health. Our findings robustly demonstrate that dietary factors exert significantly stronger influence than genetic ancestry on microbial diversity (PERMANOVA R²=0.10 vs. 0.07, p<0.05), community structure, and taxonomic composition – challenging established ethnicity-centric microbiota paradigms[19] . This dietary dominance aligns with global patterns observed across agricultural populations, where subsistence diets drive convergent microbial evolution independent of geographical separation[8; 11]. The Yi microbiome exhibits specialized enzymatic machinery for processing resistant starches, evidenced by Prevotellaceae dominance (log₂FC=1.82, padj=0.028) and Succinivibrionaceae enrichment (log₂FC=2.15, padj=0.013), corresponding to their buckwheat-rich diet (58.5±12.3% vs. Han 3.3±1.8%, p<0.001). Phylogenetic analysis revealed these taxa form a monophyletic clade (bootstrap=92%), suggesting co-adaptation to carbohydrate-rich diets[20]. The strong co-occurrence between Prevotellaceae and Succinivibrionaceae (Spearman ρ>0.6) reflects syntrophic metabolic coupling[21], where Prevotellaceae degrades complex starch into oligosaccharides for Succinivibrionaceae conversion. This consortium enhances energy harvest efficiency but associates with BMI elevation (ß=+41.8) during nutritional transition – a phenomenon termed the "Asian obesity paradox" observed in rapidly modernizing populations[9].

Conversely, Han microbiomes showed enriched Bacteroidaceae (log₂FC=1.24, padj=0.041), aligned with higher animal fat intake (23.6% vs. Yi 1.8%, p<0.001). Functional prediction indicated enhanced bile acid metabolism (terpenoid pathway enrichment 1.06-fold, p=0.006), potentially explaining paradoxical adiposity suppression (ß=-29.2) through anti-inflammatory secondary bile acid production[22]. Tripartite network analysis revealed (Fig. 4B) further delineated three core diet-microbe-function axes: (1) the buckwheat-Prevotellaceae association (r=0.42) with enhanced amino acid metabolism (r=0.41); (2) bacon consumption linkages to Enterobacteriaceae; and (3) Bacteroidaceae’s dual correlations with glycan biosynthesis (r=0.66) and suppressed terpenoid metabolism (r=−0.58). The inverse relationship between glycan biosynthesis and terpenoid metabolism in Bacteroidaceae likely reflects fundamental microbial resource allocation principles. Bacteria often prioritize metabolic pathways based on substrate availability, and the observed pattern suggests potential trade-offs where glycan utilization may divert resources from bile acid transformation. Similarly, the bacon-Enterobacteriaceae linkage aligns with established ecological patterns where processed meats create intestinal environments favoring these bacterial groups.

These statistically significant correlations represent important observational relationships, though mechanistic causation requires experimental validation. Bacon effects may be confounded by preservatives or cooking methods, and the Bacteroidaceae metabolic trade-off hypothesis warrants verification through gene expression profiling. Future studies should quantify bile acid profiles across dietary groups and conduct controlled interventions to isolate specific food effects.

The 12% higher functional redundancy in Yi microbiomes (p<0.0001) with narrower distribution (CV=10.7% vs. Han 16.9%) creates ecological resilience against dietary perturbations[14], though direct evidence from perturbation experiments is required. This stability, mediated by Prevotellaceae-Succinivibrionaceae networks, facilitates metabolic channeling during nutrient scarcity[23] and mirrors adaptations in other high-fiber-consuming populations like the Hadza[11]. However, urbanization-associated erosion of this redundancy correlates with loss of keystone hydrogenotrophic methanogens[24], reducing metabolic flexibility and short-chain fatty acid production capacity [9]. This extends the biodiversity-ecosystem functioning framework[25] to human microbiomes, suggesting traditional diets maintain functional plasticity lost in industrialized populations [26].

The buckwheat intervention enriched *Prevotellaceae*, which are known β-glucan degraders[27]. Their expansion correlated with acetate accumulation (*r*=0.38, *p*=0.01), potentially activating FFAR2 signaling—a pathway shown to stimulate GLP-1 secretion in preclinical models[28]. Current evidence supports buckwheat as a functional food for metabolic health, with observed benefits in glucose regulation and promising microbiota-mediated mechanisms that warrant deeper investigation.

Structural equation modeling suggested a pathway linking ethnicity → buckwheat intake → microbiota → cardiometabolic traits. Concurrently, *Oscillospiraceae* and *Ruminococcaceae* abundances positively correlated with HDL levels (*r*=0.221 and 0.198)( Supplementary 1). Given their butyrate-producing potential and evidence that butyrate activates LXRα-dependent reverse cholesterol transport in mice[29], these taxa may represent novel targets for atherosclerosis intervention, pending mechanistic confirmation."

Our findings reveal that traditional Yi diets associate with gut microbiome features through quantifiable evolutionary and ecological patterns. Phylogenetic analysis indicates that Yi-enriched Prevotellaceae and Succinivibrionaceae form a monophyletic clade (bootstrap=92%), implying potential shared evolutionary trajectories under high-resistant starch selection. This dietary association manifests functionally through enhanced glycan metabolism (1.17-fold, p=0.005) and ecologically via elevated functional redundancy (12% higher, p<0.0001) – a metric potentially conferring stability advantages. The buckwheat-Prevotellaceae correlation (r=0.42, p=1.1×10⁻⁵) exemplifies diet-driven microbial niche partitioning, while Oscillospiraceae-HDL covariation (r=0.221) implies statistical association requiring mechanistic validation.

Methodologically, the integration of phylogenetics, networks, and functional metrics provides a multidimensional analytical framework. Observed convergence of carbohydrate-utilizing taxa into phylogenetically conserved consortia, coupled with network-structured functional capacity, offers a model for investigating diet-microbiome relationships.

Future longitudinal studies should resolve strain-level dynamics within Prevotellaceae-Succinivibrionaceae networks during dietary transitions, explicitly testing whether the observed functional redundancy difference confers measurable ecosystem stability. Concurrent tracking of microbial glycan metabolism and HDL covariation is needed to establish temporal precedence in any diet-microbiota-host interactions.

## Materials and methods

### Population

The study was carried out in Liangshan district, located in Southwest China, from 01/10/2021 to 30/11/2022. Ethnic classification was determined by residential area and official registration: Yi participants resided in predominantly Yi communities of Liangshan with Yi ethnicity registered in the national household registry; Han participants resided in Han-majority areas with Han ethnicity registration. Eligible participants also met the following criteria: (i) no antibiotic or probiotic use in the recent three months ; (ii) no gastrointestinal disease( for example, enteritis, enterorrhagia,Diarrhea); (iii) no medical history of chronic diseases or metabolic disorders(for example, hypertension, cardiovascular disease, T2DM). A total of 100 donors were included in the study , consisting of 50 Yi and 50 Han participants.

### Demographic and Clinical Data Collection

Anthropometric measurements (height, weigh) followed standardized WHO STEPS protocols. Age, sex, ethnicity, and food type (Cooked rice, Cooked buckwheat, Bacon(raw), Freshmeat (raw) , Vegetables(raw), Fruits (raw) ) were recorded through questionnaires. Blood samples were collected after overnight fasting for biochemical analyses including lipid profiles (HDL, LDL, triglycerides) and glucose. Body mass index (BMI) was calculated as weight (kg) divided by height squared (m²).

### Dietary Assessment

Dietary intake was assessed through three non-consecutive 24-hour recalls[30] (including one weekend day) using standardized portion-size photographs and household measures, with daily intake (g/day) calculated via the Chinese Food Composition Tables; validation included a subset (n=30) completing 3-day weighed food records and plasma biomarkers analysis (alkylresorcinols for whole grains, β-carotene for vegetables), focusing on five food categories: rice, buckwheat, bacon, fresh meat, and vegetables.

### Stool sample collection

We optimized DNA extraction from human fecal samples by incorporating a lysozyme pretreatment (10 mg/mL, 37°C, 30 min) and mechanical lysis with 0.1 mm zirconia beads to enhance Gram-positive bacterial recovery, followed by strict quality control (A260/A280 >1.8, gel verification). For 16S rRNA amplification, we used adapter-tailed primers 338F/806R with reduced cycle numbers (20 cycles) and adjusted annealing temperature (50°C) to minimize bias in low-biomass samples, followed by size-selection (550±50 bp) and paired-end sequencing (2×250 bp) with batch-specific negative controls (standard protocols provided in Supplementary Materials)

### Microbiota Diversity and Composition Analysis

Microbial α-diversity was quantified using the Shannon index, while β-diversity was assessed through Bray–Curtis dissimilarity and visualized via principal coordinates analysis (PCoA). Differences in β-diversity between ethnic groups were tested using permutational multivariate analysis of variance (PERMANOVA)[31] with 999 permutations in the vegan R package. Critically, all analyses incorporated statistical adjustments for observed demographic differences: DESeq2 models[32] explicitly included age, sex, and BMI as covariates, while PERMANOVA implemented covariate stratification. Differential abundance of bacterial taxa was subsequently analyzed using DESeq2 with negative binomial generalized linear models. Finally, microbial co-occurrence networks were constructed through SparCC correlations[33] (absolute value >0.3 threshold) with 100 bootstrap iterations, followed by false discovery rate (FDR) correction, and visualized in Gephi 0.9.7 using the Fruchterman–Reingold layout algorithm.

### Functional Prediction and Tripartite Network Analysis

Functional potential of the microbiota was predicted using PICRUSt2[34] (Phylogenetic Investigation of Communities by Reconstruction of Unobserved States) with default parameters. Predicted Kyoto Encyclopedia of Genes and Genomes (KEGG) pathways were normalized by cumulative sum scaling (CSS) in the metagenomeSeq R package[35]. Differential pathway abundance between groups was tested using Welch’s t-test with Benjamini–Hochberg FDR correction in STAMP v2.1.3. <u>Covariate adjustment</u> was implemented in all functional comparisons. Tripartite networks integrating dietary factors, microbial families, and KEGG pathways were constructed based on Spearman correlations (absolute r > 0.3, FDR-adjusted p < 0.05). Gaussian graphical models[36] were used to infer conditional dependencies, and networks were visualized using the igraph R package.

### Functional Redundancy Analysis

Functional redundancy was quantified using the framework proposed by Louca et al[37], which accounts for both functional and phylogenetic diversity. The redundancy index was calculated as the ratio of ASV richness to the exponential of the Shannon diversity of KEGG pathways, multiplied by the ratio of observed phylogenetic diversity to maximum possible phylogenetic diversity. Differences between ethnic groups were tested using the Wilcoxon rank-sum test.

### Structural Equation Modeling

We used structural equation modeling to test the pathway ethnicity → buckwheat intake → gut microbiota → cardiometabolic traits. Path coefficients were estimated via maximum likelihood (adjusting for age, sex, BMI), with key microbiota families selected based on BMI associations (Fig5B). Model fit met standard thresholds (CFI>0.95, RMSEA<0.06, SRMR<0.08).

### Statistical Analysis

All statistical analyses were conducted in R version 4.2.1. Continuous variables were compared using Wilcoxon rank-sum tests or Student’s t-tests as appropriate for data distribution, while categorical variables were assessed with χ² tests. Multiple testing correction was applied using the Benjamini-Hochberg false discovery rate (FDR) method. To ensure robustness of findings, sensitivity analyses included: (i) subsampling to address covariate imbalances, (ii) leave-one-out cross-validation, and (iii) batch effect correction via ComBat-seq. Statistical significance was defined as p < 0.05 or FDR-adjusted q < 0.05 for all analyses.

## Acknowledgments

We thank all the staff who contributed to this study.

The authors have declared that no competing interests exist.

## Impact Statement

This study demonstrates that traditional dietary practices, rather than genetic ancestry, primarily drive gut microbiome divergence in cohabiting ethnic groups. We identify buckwheat consumption as a key modulator of Prevotellaceae-dominated consortia with enhanced metabolic functionality and ecological resilience, offering actionable strategies for microbiota-targeted nutritional interventions in multi-ethnic communities.

## Ethical Approval Statement

This study was conducted in accordance with the Declaration of Helsinki and was approved by the Ethics Committee of First People’s Hospital of Liangshan Yi Autonomous Prefecture, Sichuan University and Huazhong University of Science and Technology. All procedures followed relevant guidelines and regulations. Written informed consent was obtained from all participants or their guardians prior to enrollment.

## Data availability

The raw sequencing reads generated in this study are publicly available in the NCBI Sequence Read Archive (SRA) under BioProject accession [PRJNA1314298](https://www.ncbi.nlm.nih.gov/bioproject/PRJNA1314298).

